# Modeling the development of *Triatoma infestans* through temperature: Estimating generation time in Bolivia

**DOI:** 10.1101/2023.08.31.555655

**Authors:** Frédéric Lardeux, Stéphanie Depickère, Rosenka Tejerina

## Abstract

**Background:** In Bolivia, controlling *Triatoma infestans*, the primary vector of Chagas disease, remains challenging in the hot regions of the country. The study aims to establish a temperature-based model of development for *T. infestans* and explore phenological factors that could partially explain the failures in vector control within these regions of high ambient temperature.

**Methods:** We employed the Briere-1 model to describe the development time from egg-hatch to adults of *T. infestans* with temperature. Given that the entire developmental cycle can exceed two years under cooler temperature conditions, direct study of this duration was not undertaken. Instead, simulation was employed. For this purpose, insect cohorts encompassing all six stages (egg, N1, N2, N3, N4, N5) were concurrently raised within temperature-controlled climate chambers. The number of days required for molting between consecutive stages was recorded. Using this recorded dataset, the development time from eggs to adults was statistically simulated for various constant temperatures. The Briere-1 model was then calibrated using the dataset from each molting phase and applied to the simulated complete development cycle. The model was then used in conjunction with field temperatures from four representative localities within Bolivia to compute development times and generation intervals. A GIS approach was also used to map development times and generation intervals in the geographical distribution range of *T. infestans*.

**Findings:** The model suggests that the minimum temperature required for the development of *T. infestans* is approximately 15°C. The temperature at which its development attains maximum efficiency is around 33°C, while the threshold for lethal temperature stands at approximately 39°C. In the warmer regions of Bolivia, *T. infestans* exhibits an almost bivoltine cycle, with the number of yearly generations (*G*) ranging from approximately 1.5 to 2.5. In contrast, within the cooler Dry Inter-Andean Valleys, its cycle becomes univoltine or even less frequent (*G*≤1). The model could potentially offer insights into the correlations between insecticide resistance and the number of yearly generations, thereby clarifying why the control of *T. infestans* in hotter regions proves more challenging to achieve.

**Interpretation:** The notion of generation time arises as a pivotal consideration in the management of *T. infestans*, especially within Bolivia’s warmer regions. In areas marked by higher temperatures, the generation time of the vector diminishes, leading to a notable increase in the population growth rate. This, in turn, accelerates the emergence of insecticide resistance, as evidenced by the findings of this current study.

**Author summary:** In Bolivia, the bug *Triatoma infestans* is the main carrier of *Trypanosoma cruzi*, the parasite responsible for Chagas disease. This study aims to understand how these bugs develop in different temperatures and how temperature affects efforts to control Chagas disease.

Experiments in controlled environments with different temperatures were carried out and results revealed that the Briere-1 model can accurately imitated how the bugs’ growth rate changes with temperature: following a sigmoid pattern, higher temperatures make the bugs grow faster up to a maximum before slowing rapidly down. Using this model, the study looked at how long it takes for a new generation of bugs to develop and therefore estimated the generation time as a function of temperature. It appeared that in warmer places, the bugs can have more than one generation in a year, which makes their population grow quickly and increases the risk of Chagas disease spreading.

The study also looked at whether the bugs’ resistance to insecticides is correlated to the generation time and it appeared that areas where bugs reproduce quickly tend to have more resistance to insecticides.

This research emphasizes how important it is to consider temperature when trying to control *T. infestans*. Indeed, in areas with higher temperatures, the bugs reproduce more quickly and show a greater tendency to develop higher levels of resistance to insecticides. This dynamic complicates efforts to control them effectively. This information holds the potential to inform the development of improved strategies to curtail the spread of Chagas disease.

## Introduction

In Bolivia, the reduviid bug *Triatoma infestans* is the primary vector of *Trypanosoma cruzi*, the causative agent of Chagas disease. Within this country, over 3.5 million individuals reside in endemic regions where the risk is prevalent. A significant portion, around 6% of the Bolivian population, is affected by the disease. Regrettably, every year, thousands of people continue to contract the disease through vectorial transmission. *T. infestans* predominantly proliferates within impoverished rural residences or adjacent structures, such as household storages, poultry houses, hutches, or cattle enclosures [1]. In these dwellings, for years, the Bolivian National Chagas Program has undertaken initiatives to diminish household infestations through the application of residual insecticides. While certain positive outcomes have emerged, including the certification of vectorial transmission interruption in select Andean regions (notably temperate areas), Chagas disease transmission remains active, particularly in the Chaco region located in the southern part of the country characterized by warmer climates [2]. The Gran Chaco ecoregion spans Argentina, Bolivia, and Paraguay, where the prevalence of *T. infestans* and household infestations is influenced by factors such as residents’ living habits, including the construction materials used (such as mud-and-thatch materials, which provide a suitable habitat for *T. infestans*), and other sociocultural practices that have contributed to the proliferation of vector populations [1]. Despite efforts to reduce vector numbers through insecticide applications, effective control over *T. infestans* has yet to be attained within this ecoregion [3]. Numerous factors have been proposed to account for this shortcoming, including: (1) political instability leading to inconsistent vector control campaigns by National Control Programs; (2) the presence of *T. infestans* populations resistant to insecticides [3, 4]; and (3) limited efficacy of standard spraying techniques and pyrethroid formulations, often not properly executed by technicians [5]. Additionally, the frequency of insecticide applications might be inadequate, as evidenced by the superior outcomes yielded by long-lasting insecticide formulations compared to their shorter-lived counterparts. This indicates that when the efficacy of an insecticide diminishes, insect populations could recover between interventions [6]. Consequently, for *T. infestans*, the frequency of generations per year, or equivalently, the generation time (*i.e.*, the period encompassing a complete developmental cycle or the average span between one generation’s birth and the next), becomes a pivotal statistic when implementing chemical control measures. For *T. infestans*, this generation time hinges on the developmental rate, itself dictated by ambient temperature, a fundamental factor for poikilothermic organisms. Phenological models, regularly employed to forecast population statistics (including generation time), are pivotal for refining pest management tactics [7]. Among these models, those driven by temperature hold substantial value in crafting pest management strategies. Temperature, as a paramount parameter in insect phenology, makes temperature-driven models—though straightforward—generally sufficient to delineate insect population dynamics in various climatic contexts. This empowers pest managers to make informed decisions.

Hence, the core objective of this study is to introduce a temperature-driven developmental model for *T. infestans*. The model serves as an initial effort to compute generation times for *T. infestans* in Bolivia and explore phenological aspects that could partly elucidate the failures of vector control, particularly in regions characterized by higher temperatures.

## Methods

### Breeding of *T. infestans* in temperature-controlled environment

The strain of *T. infestans* utilized in all experiments was the CIPEIN strain [8]. Batches of insects that had recently entered a developmental stage (egg, Nymph 1 (N1), N2, N3, N4, or N5) were placed within climatic chambers (Binder KBWF 720, Tuttlingen, Germany) set at operational temperatures of 20, 25, 27, 30, 35, and 37°C (+/-0.1°C), with relative humidity approximately ranging between 40-60%. The total number of processed insects (7988 in total, divided among the 6 stages and the 6 temperatures) is presented in Table 1.

**Table 1.**
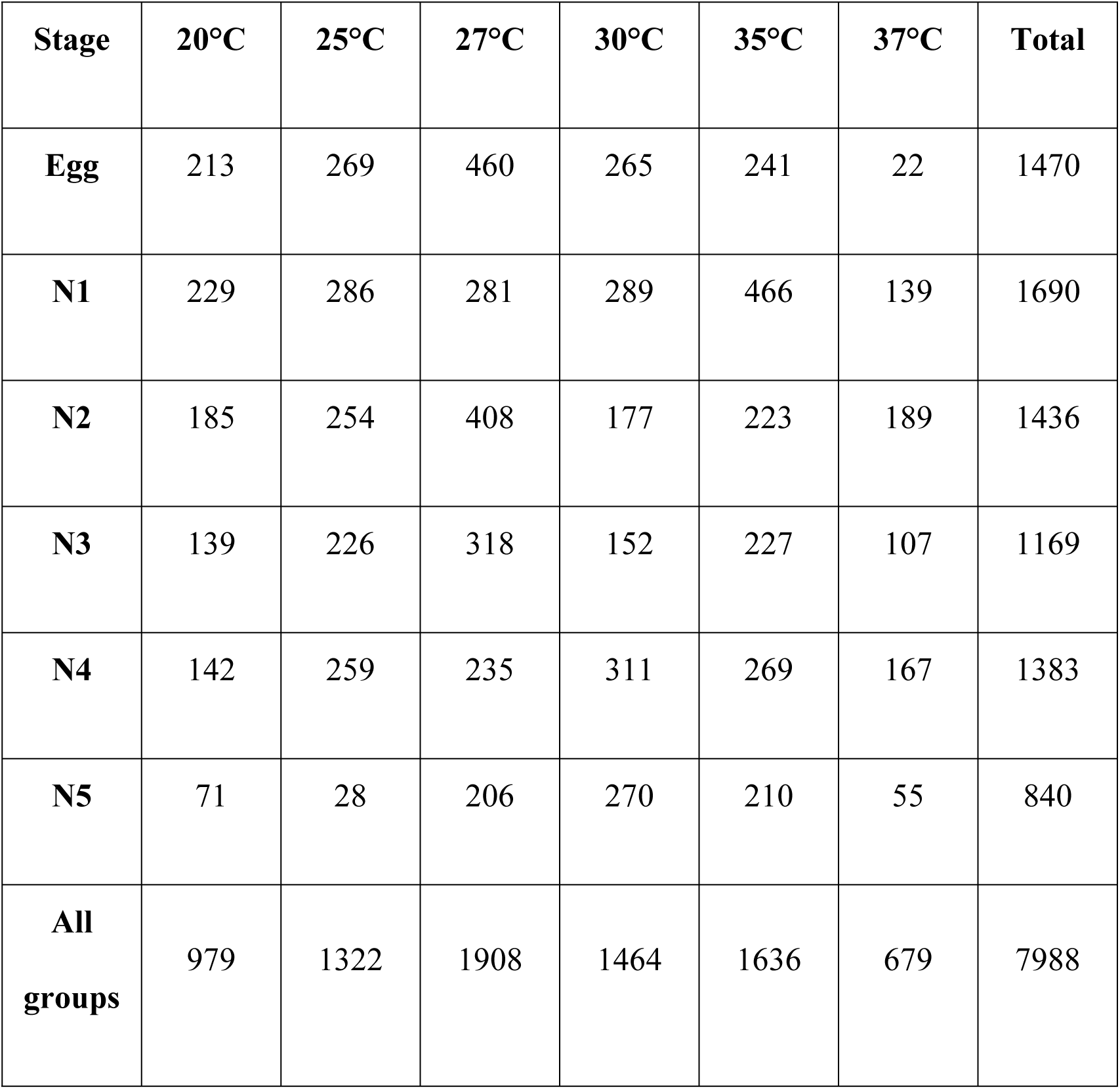
Number of processed individuals at each working temperature.

Eggs were carefully positioned within Petri dishes, while nymphs were accommodated in plastic containers equipped with accordion-folded craft paper (2-liter plastic containers for N1, N2, N3 stages, and 5-liter plastic containers for N3, N4 and N5 stages). Although attempts were made at 16°C and 40°C, no development was observed (insects did not undergo molting). Furthermore, at 40°C, insects experienced rapid mortality. Under other different temperatures, nymphal stages were permitted to feed on chicken blood once a week. Daily counts were performed on insects as they transitioned to the subsequent stage. The duration of time each insect spent in a particular stage was meticulously recorded, along with the rearing temperature, to create the working dataset (S1 Table).

### Modelling *T. infestans* development with temperature

The development rate is quantified as the portion of the developmental period occurring within a given calendar time frame. By definition, development is considered accomplished when the accumulated development rates sum up to 1 (indicating 100% completion). Hence, the development rate is gauged as the reciprocal of the number of time units necessary for completing the developmental process. This measurement can encompass the entire ontogenesis (from egg to adult in the context of *T. infestans*) or focus on a specific stage, such as, for example, the time required for N3 to N4 molting. In this study, the chosen time unit for assessment was the day. Mathematically, the accumulated development rates for complete development can be expressed as [9]:

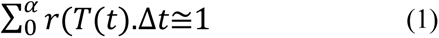

Where *r(T(t))* represents the temperature-dependent development rate at time *t*. *T* is the temperature, and α symbolizes the total development duration (*i.e.,* the parameter to be estimated). The intervals of constant temperature are denoted as *Δt*, and in this context, they correspond to daily periods, as described in this paper.

For each point in time *t* throughout the day, corresponds a specific temperature *T(t),* alongside an associated rate of development *r(T(t)*). The model hinges on the requirement of the function *r(T),* representing the development rate *r* at the given temperature *T*. Various functions have been proposed [10]. In the current study, the Brière-1 function [11] was adopted, having exhibited the lowest Akaike Information Criterion (AIC) scores when compared to other phenological models included in the R “devRate” package [12] with the present dataset.

The Brière-1 model expresses the development rate *r(T)* at temperature *T* as:

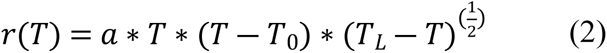

Where *a* is a constant, *T_0_* is the lower temperature threshold (which represents the critical thermal minimum where development rates slightly exceed 0) and *T_L_* is the upper temperature threshold (above which rates are 0).

Another crucial parameter is *Topt*, denoting the optimal temperature at which the development rate achieves its peak. All these parameters can be estimated by the R-devRate package.

The Brière-1 model was adjusted to each of the six developmental phases of *T. infestans*: [egg to N1], [N1 to N2], [N2 to N3], [N3 to N4], [N4 to N5], and [N5 to adult]. The complete cycle (eggs to adults), which can span over two years or more under lower temperatures, wasn’t directly studied in the climatic chambers. However, it was simulated using observed data and the @RISK software (Palisade Co, Ithaca, NY, USA) as follows:

Using the observed development rate histograms at a specific operational temperature, an initial development rate (from egg to N1) was assigned to each of the 100 000 simulated individuals. This assignment was done with a probability equivalent to the rate’s frequency in the histogram of rates of this first cohort. The software then generated a resultant histogram of development rates for the egg to N1 phase. These computational steps were iteratively applied to the resulting histogram to generate simulated histograms for subsequent developmental phases (N1 to N2, N2 to N3, and so forth), culminating in the final developmental phase (N5 to adult) (S2. Table). The resultant composite histogram depicted the distribution of development rates for the entire cycle at a given temperature. This process was replicated for all six operational temperatures, and the Brière-1 model was then adjusted to these simulated datasets.

### Cycle duration and generation time

Four distinct localities within the geographical range of *T. infestans* were selected for study. Two in the Chaco region (Yacuiba and Camiri) where temperatures are warmer, and two in the dry inter-Andean valleys of Bolivia (Cochabamba and Mizque) where temperatures are cooler (Table 2). Mean daily temperature data for these localities were sourced from SENAMHI, the meteorological service of Bolivia (http://www.senamhi.gob.bo/) (S3 Table).

**Table 2.**
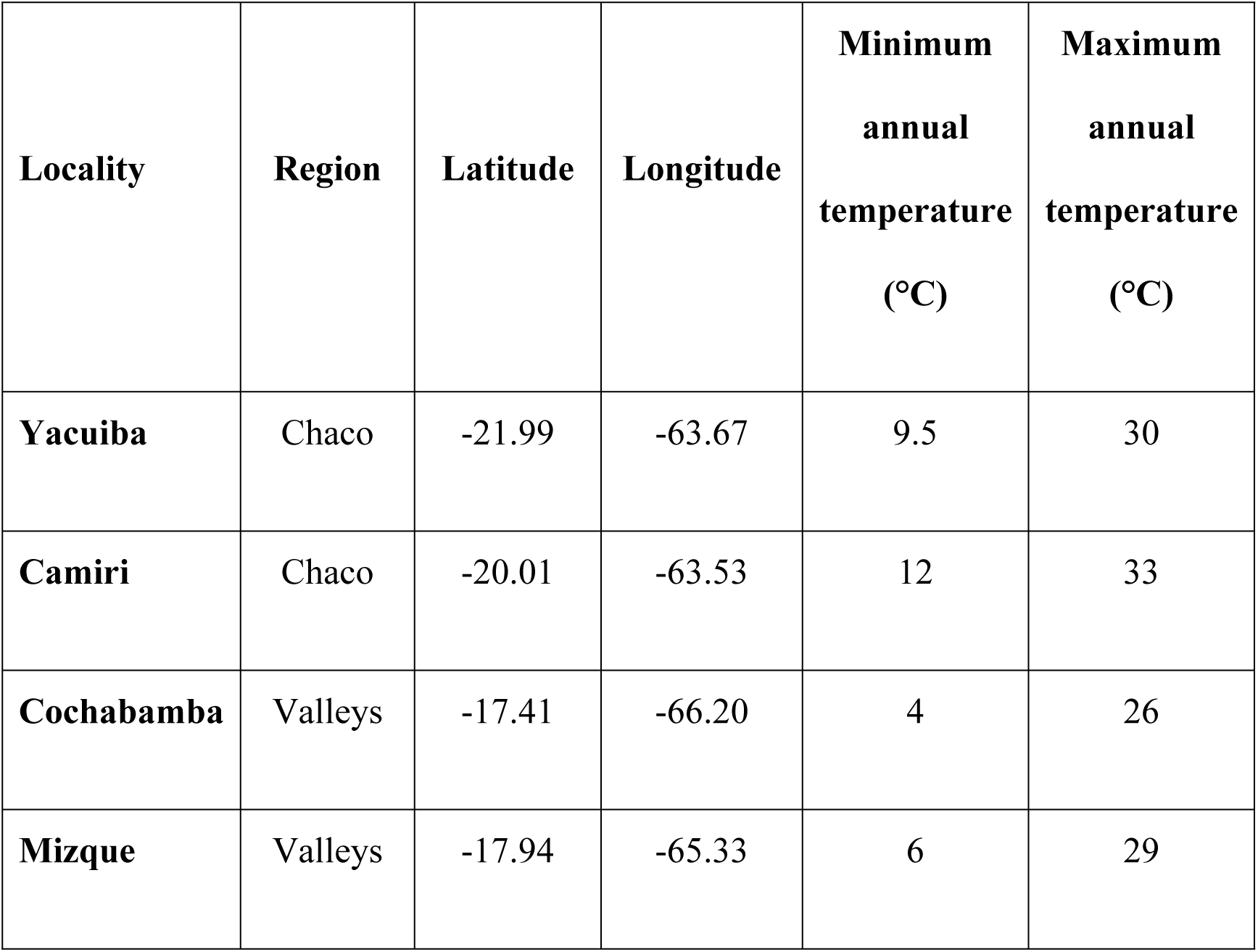
Bolivian localities used to compute development rates and generation times of T. infestans.

Using the mean daily temperatures from January 2012 to July 2017, coupled with the Brière-1 model fitted for the complete *T. infestans* life cycle (from egg to adult), the time duration of the cycle was calculated for an individual born on day *t*. This computation was performed across this time frame (*<u>i.e,</u>* almost 5.5 years) and within each of the selected localities, employing equation (1). Moreover, the number of generations during the 2012-2017 period for each locality was determined utilizing the devRate package. This was achieved by assuming an additional 15-day span between adult emergence and the initial oviposition, while simulating the population growth of 50 individuals. The devRate package doesn’t encompass population variables like mortality; as a result, only the timeframe of each developmental stage is simulated. Within this simulation, the entire sequence of developmental stages, ranging from egg to N1, then N2, and onward to adults that lay eggs, is undergone by each of the 50 individuals. Subsequently, in the next generation, the same cohort of 50 individuals progresses through successive stages anew, continuing this pattern.

Based on this simulation, graphical representations illustrating the progression of each developmental phase (egg-N1, N1-N2, and so forth) over time were generated for each locality throughout the specified period.

The devRate package was also used to compute the number of generations of *T. infestans* per year using WorldClim data as follow: Average monthly temperatures for Bolivia spanning the period 1972-2000 were extracted from the tavg30s file (30-minute resolution), accessible at https://www.worldclim.org/data/worldclim21.html. For each pixel within the WorldClim dataset, development rates were computed for every month across the year, and then aggregated to derive the annual development rate. The computed annual development rate corresponds to the mean number of generations per year. The graphical representation of the mean number of generations per year was carried out with QGIS v.3.10 (https://www.qgis.org/en/site/), within the approximate potential geographical distribution range of *T. infestans* in Bolivia. [13]

The rate of appearance of insecticide resistance is directly contingent upon the generation time, while other contributing factors (notably the selection pressure as influenced by the intensity of control efforts), exhibit more of a logarithmic impact [14]. This study scrutinizes the pivotal role of generation time in the resistance phenomenon, leveraging published resistance data of *T. infestans* toward deltamethrin, an insecticide deployed by the Bolivian National Chagas Program in its control operations. The dataset comprises computed resistance ratios (RR) at 50% lethal concentrations (LD50) for deltamethrin, contrasting the CIPEIN susceptible reference strain against strains gathered from 21 distinct localities, each with known GPS coordinates [3]. For these localities, the number of generations per year was derived from the pixel values of the GIS layer corresponding to each GPS point of the localities. A graphical representation was generated to elucidate the correlation between resistance level (RR) and the average number of generations within each locality.

## Results

### Effect of temperature on development and cycle duration

The outcomes of constant-temperature rearing experiments are succinctly presented in Table 3. The average duration of the complete cycle was ≈ 407 days at 20°C (range: [274-563]), and ≈ 99 days (range: [74-151]) at 35°C, a temperature closely aligned with the optimal temperature computed by the Briere-1 model. The parameters of the model for each stage and the entire cycle, in conjunction with the computation of *Topt*, are comprehensively summarized in Table 4.

**Table 3.**
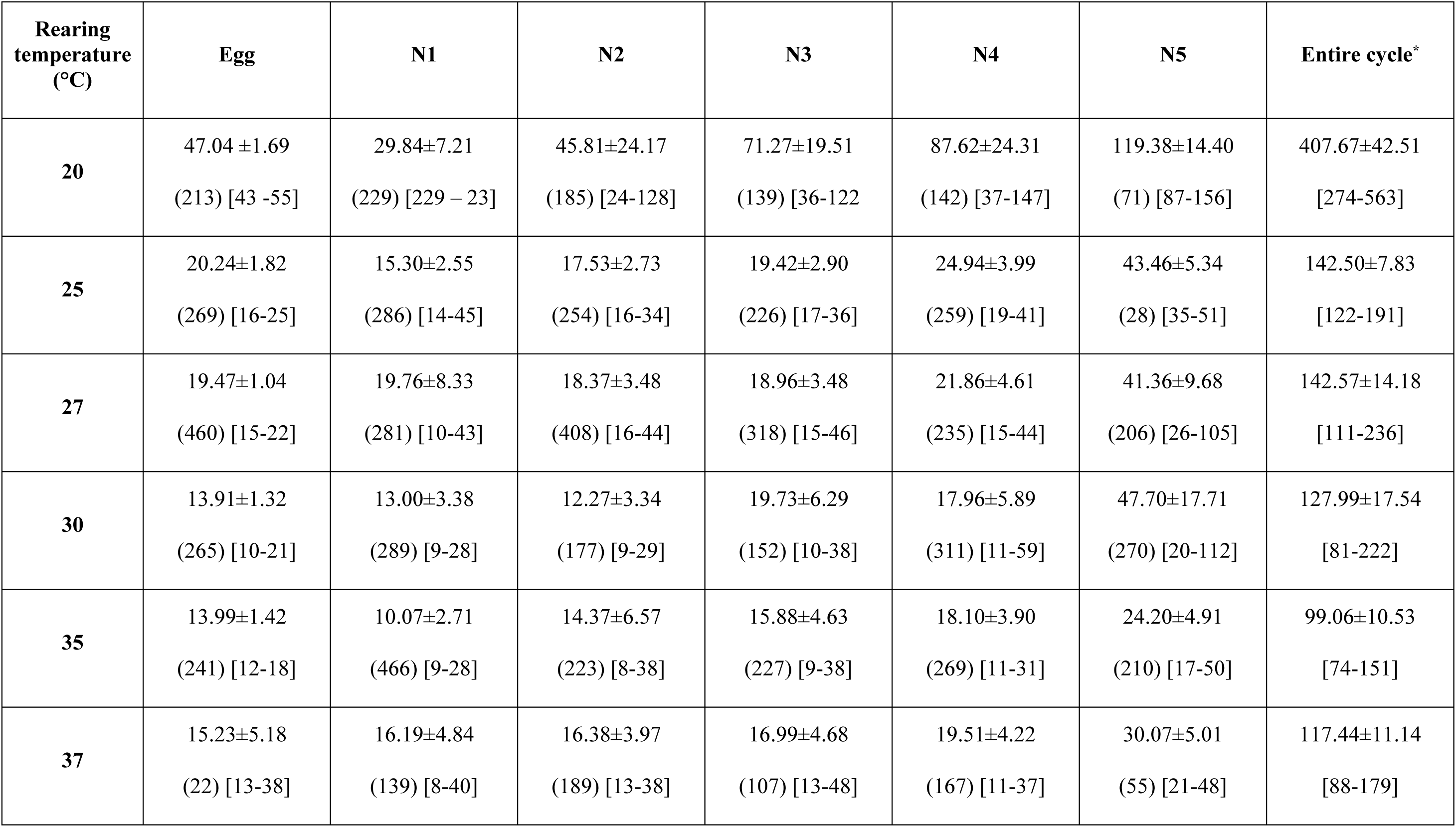
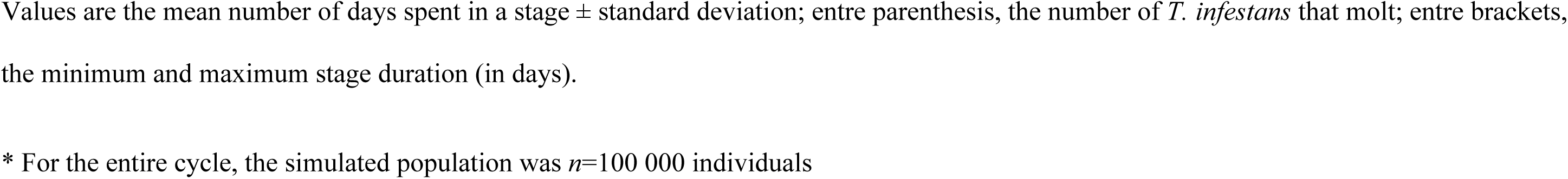
Cycle duration of *T. infestans* at various constant temperatures.

| **Rearing temperature (°C)** | **Egg** | **N1** | **N2** | **N3** | **N4** | **N5** | **Entire cycle\*** |
| --- | --- | --- | --- | --- | --- | --- | --- |
| **20** | 47.04 ±1.69<br>(213) [43 -55] | 29.84±7.21<br>(229) [229 – 23] | 45.81±24.17<br>(185) [24-128] | 71.27±19.51<br>(139) [36-122] | 87.62±24.31<br>(142) [37-147] | 119.38±14.40<br>(71) [87-156] | 407.67±42.51<br>[274-563] |
| **25** | 20.24±1.82<br>(269) [16-25] | 15.30±2.55<br>(286) [14-45] | 17.53±2.73<br>(254) [16-34] | 19.42±2.90<br>(226) [17-36] | 24.94±3.99<br>(259) [19-41] | 43.46±5.34<br>(28) [35-51] | 142.50±7.83<br>[122-191] |
| **27** | 19.47±1.04<br>(460) [15-22] | 19.76±8.33<br>(281) [10-43] | 18.37±3.48<br>(408) [16-44] | 18.96±3.48<br>(318) [15-46] | 21.86±4.61<br>(235) [15-44] | 41.36±9.68<br>(206) [26-105] | 142.57±14.18<br>[111-236] |
| **30** | 13.91±1.32<br>(265) [10-21] | 13.00±3.38<br>(289) [9-28] | 12.27±3.34<br>(177) [9-29] | 19.73±6.29<br>(152) [10-38] | 17.96±5.89<br>(311) [11-59] | 47.70±17.71<br>(270) [20-112] | 127.99±17.54<br>[81-222] |
| **35** | 13.99±1.42<br>(241) [12-18] | 10.07±2.71<br>(466) [9-28] | 14.37±6.57<br>(223) [8-38] | 15.88±4.63<br>(227) [9-38] | 18.10±3.90<br>(269) [11-31] | 24.20±4.91<br>(210) [17-50] | 99.06±10.53<br>[74-151] |
| **37** | 15.23±5.18<br>(22) [13-38] | 16.19±4.84<br>(139) [8-40] | 16.38±3.97<br>(189) [13-38] | 16.99±4.68<br>(107) [13-48] | 19.51±4.22<br>(167) [11-37] | 30.07±5.01<br>(55) [21-48] | 117.44±11.14<br>[88-179] |

Values are the mean number of days spent in a stage ± standard deviation; entre parenthesis, the number of *T. infestans* that molt; entre brackets, the minimum and maximum stage duration (in days).
\* For the entire cycle, the simulated population was $n=100\ 000$ individuals

**Table 4.** Parameters estimates of Brière-1 model, for each instar and the entire cycle of T. infestans.

| Stage | $a^*$ | $T_0$ | $T_L$ | $T_{opt}$ | Thermal<br>breath |
| --- | --- | --- | --- | --- | --- |
| **Egg** | 4.882 | 15.12 | 39.69 | 33.7 | 23.3–39.1 |
| **N1** | 5.797 | 14.18 | 39.89 | 33.7 | 22.9-39.2 |
| **N2** | 5.893 | 15.36 | 38.88 | 33.1 | 23.1-38.3 |
| **N3** | 4.429 | 14.42 | 39.66 | 33.6 | 22.9-39.0 |
| **N4** | 4.631 | 16.01 | 39.00 | 33.3 | 23.5-38.4 |
| **N5** | 2.978 | 17.68 | 40.14 | 34.5 | 24.8-39.6 |
| **Entire**<br>**cycle** | 0.720 | 15.62 | 39.32 | 33.5 | 23.4-38.7 |

Except for $a$ (dimensionless), the remaining parameters are temperatures expressed in °C.
\*The $a$ parameter value should be multiplied by $10^{-5}$

Numeric findings for the simulated complete cycle unfolded as follows: at 20°C, a mean development rate (*r*) of 0.00248 (Standard Deviation: 0.000262); at 25°C, *r* = 0.007050 (SD: 0.000375); at 27°C, *r* = 0.007079 (SD: 0.000684); at 30°C, *r* = 0.007957 (SD: 0.001046); at 35°C, *r* = 0.010219 (SD: 0.001036); and at 37°C, *r* = 0.008666 (SD: 0.000658).

The graph illustrating the Briere-1 model for this entire cycle showcased a prototypical development rate curve, evincing an almost linear relationship at moderate temperatures, which then transitions into a non-linear relationship at higher temperatures (Fig. 1). Across the complete cycle, the model suggested an optimal development rate around 33.5°C, akin to the optimal temperature ranges for the other stages (ranging from 33.1°C to 34.5°C). Moreover, the thermal breadth for the entire cycle, indicative of the temperature span where the development rate is at least 50% of the optimal rate, spanned from 23.4°C to 38.7°C, aligning with the ranges observed for the other developmental stages (Table 4).

**Fig. 1.**
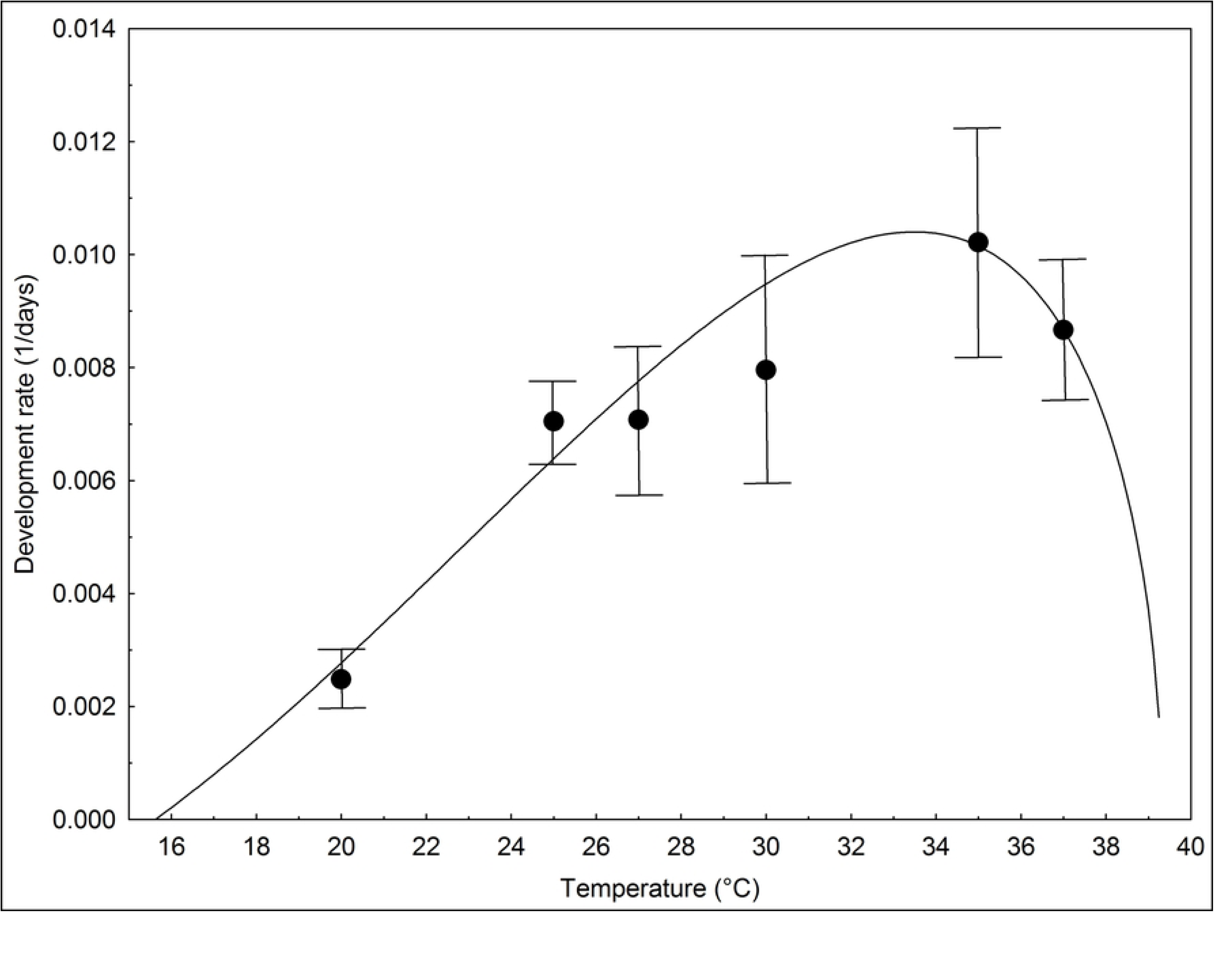
Brière-1 model adjusted to the entire development cycle of *T. infestans.* Vertical bars indicate 95% confidence interval of development rates

### Number of generations per year

For the four chosen representative localities, an overview of the average cycle duration and the number of generations per year for *T. infestans* generations is consolidated in Table 5.

**Table 5.** Mean cycle duration of *T. infestans* in the four representative localities of Bolivia, computed during the period 2012-2016.

| Localidad | Region | Annual mean temperature (°C) | Mean cycle duration | Minimum cycle duration | Maximum cycle duration | Mean number of generations per year |
| --- | --- | --- | --- | --- | --- | --- |
| Yacuiba | Chaco | 21.5 | 140.9<br>(44.4) | 83 | 227 | 1.61 |
| Camiri | Chaco | 22.9 | 130.8<br>(39.0) | 83 | 213 | 1.73 |
| Cochabamba | Valleys | 17.0 | 264.7<br>(45.8) | 163 | 338 | 0.86 |
| Mizque | Valleys | 18.9 | 205.1<br>(40.3) | 131 | 287 | 1.09 |

The annual mean temperature is in °C; the cycle durations (mean, minimum and maximum) are in days and entre parenthesis is the standard deviation of the mean.

In Camiri, situated in the hot Chaco region, the mean cycle duration was approximately 131 days, encompassing 1.73 generations annually. Conversely, in Cochabamba, nestled within the Valleys region with its cooler temperatures, the cycle stretches twice as long, covering 265 days, while the mean number of generations diminishes to half (0.86).

The entire cycle’s temporal duration across these four locales is depicted in Fig. 2. This graph illustrates the cycle duration (in days) for an individual *T. infestans* born at time *t,* plotted along the X-axis between January 2012 and December 2016. The undulating patterns observed in Fig. 2 mirror the seasonal temperature fluctuations distinctive to each region, and it’s evident that the development cycle of *T. infestans* in the Chaco region outpaces that in the Valleys region, consistently being shorter.

**Fig. 2.**
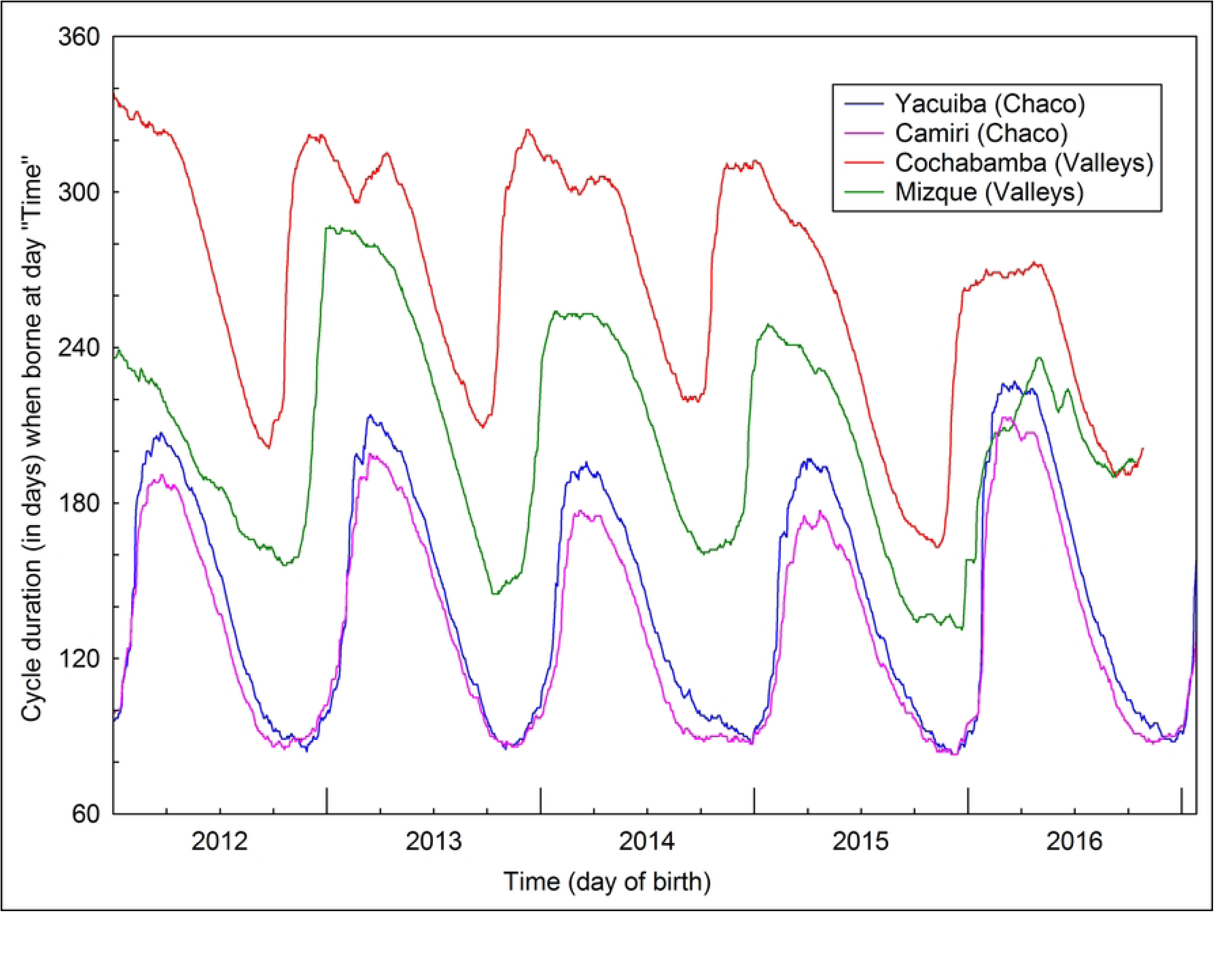
Cycle duration of *T. infestans* computed between 2012 and 2016 in four representative localities of Bolivia.

Figs 3 and 4 display the simulations of life cycles for 50 *T. infestans* in Yacuiba (Chaco region) and Cochabamba (Valleys region) respectively, executed by the devRate package. These figures depict the diverse generations of *T. infestans* that evolved within the 2012-2017 timeframe, along with the distribution of individuals at each stage of development. During this temporal span, *T. infestans* in Yacuiba (hot area) underwent nine generations of development (*i.e.*, almost 2 per year) (Fig. 3), whereas in Cochabamba (cooler area), only five generations emerged (*i.e*., one per year) (Fig. 4).

**Fig. 3.**
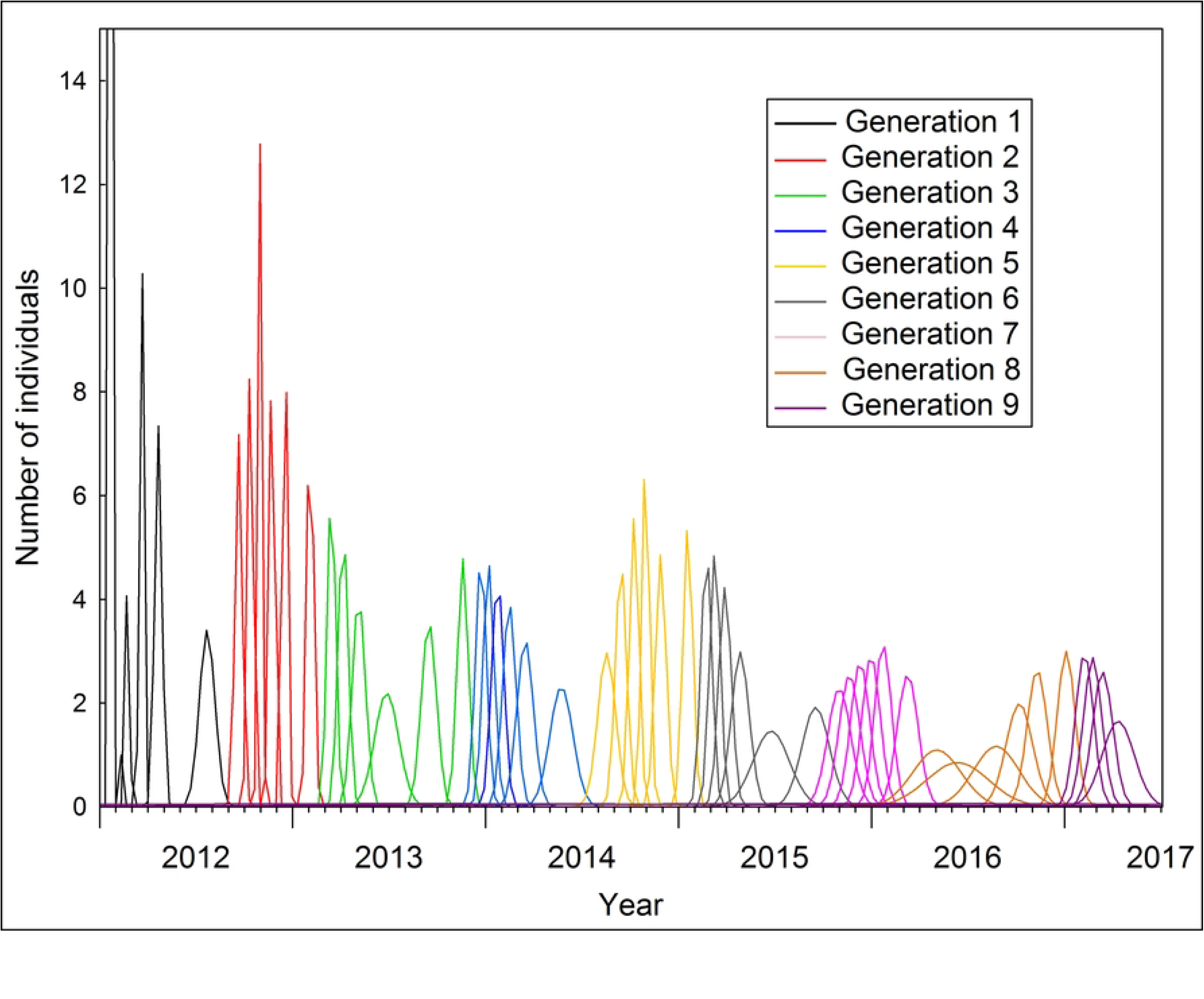
Simulation of 50 *T. infestans* development cycles during the period 2012-2017 in Yacuiba (Chaco region). Generations are identified by distinct colors and in each generation, a development stage is represented by a peak (from left to right: egg, N1, N2, N3, N4, N5).

**Fig. 4.**
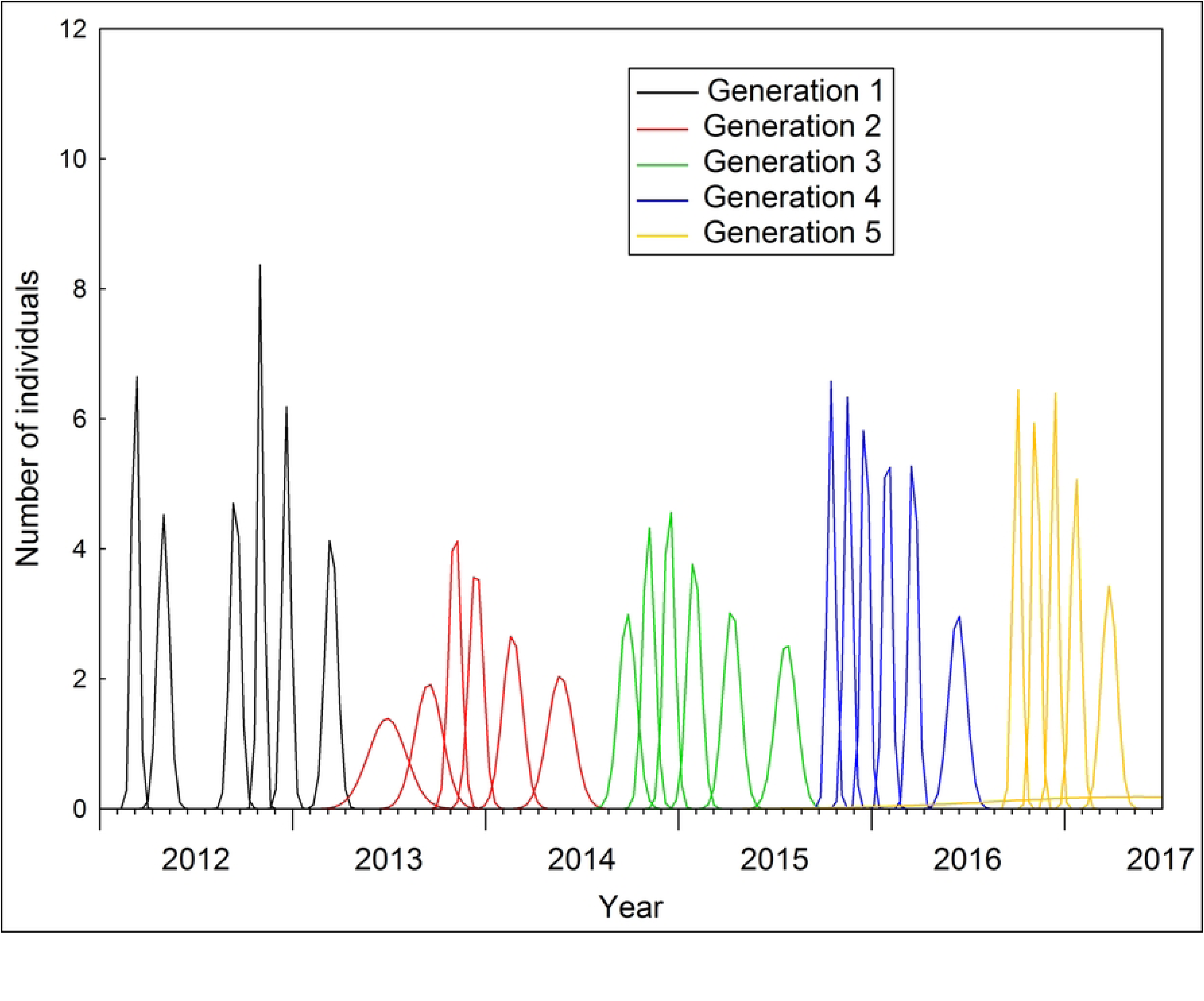
Simulation of 50 *T. infestans* development cycles during the period 2012-2017 in Cochabamba (Valleys region). Generations are identified by distinct colors and in each generation, a development stage is represented by a peak (from left to right: egg, N1, N2, N3, N4, N5).

Fig. 5 portrays the cartography detailing the number of generations per year across the conceivable geographical distribution range of *T. infestans* in Bolivia. The highest number of generations, reaching 2.43, was evident in the hottest regions like the eastern part of the Santa Cruz Department and the Chaco region. In contrast, the lowest value, registering at 0.0012, was observed in the coldest zones found within the Andes. However, while the potential distribution range of the species might suggest the presence of *T. infestans* in these cold areas, this notably low value equates to a cycle completion of 83 years – a biologically implausible span.

**Fig. 5.**
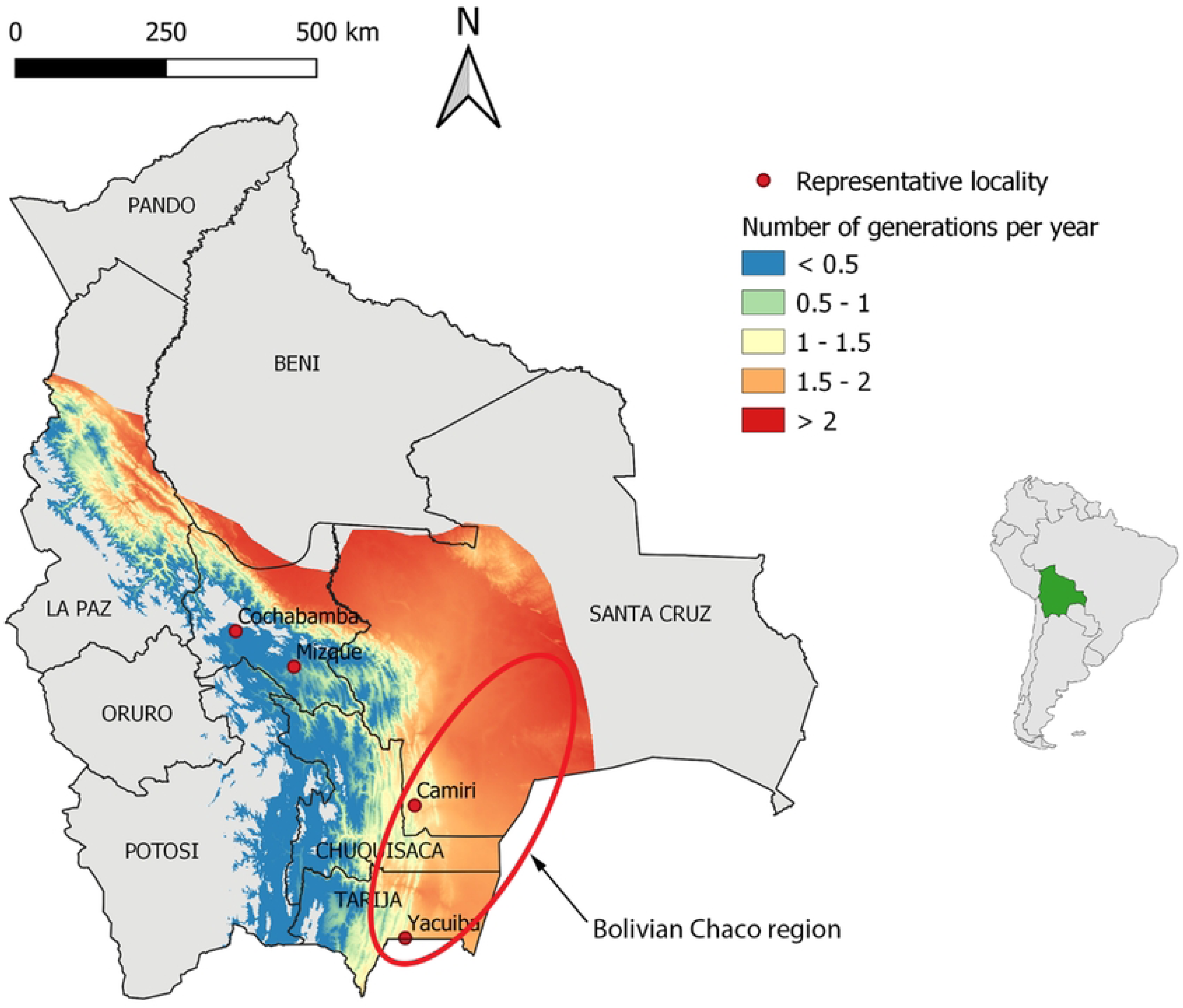
Number of generations per year for *Triatoma infestans* in its geographical distribution range in Bolivia.

In Fig. 5, the calculated number of generations per year for each pixel at the level of the four representative localities were in accordance with the outcomes listed in Table 5 and obtained through the Briere-1 model and alternative computations. Specifically, for the Valleys, the pixel values attributed to Cochabamba and Mizque were 0.57 and 0.92, respectively. In the Chaco region, the corresponding values for Camiri and Yacuiba were 1.80 and 1.73, respectively.

The depicted map in Fig. 5 delineated that within Bolivia’s lowlands – encompassing the Santa Cruz Department, the Chaco region, as well as the low-lying segments of the La Paz and Cochabamba Departments – where temperatures were comparatively warmer, the annual number of generations hovered around 1.5 or more. Across much of this expanse, the count surged above 2, even up to the peak of 2.43. This stands in stark contrast to the Andean Valleys region – characterized by higher altitudes and cooler climates – where the annual number of generations dipped below 1, approaching or even falling beneath 0.5.

The average number of off-spring that a female produce during its lifetime is the net reproductive rate (*R_0_*). For *T. infestans*, a rough estimate for *R_0_* is 150. [15] The rate of increase of a population (*r*, or Malthusian parameter) is dependent on *R_0_* and the generation time *T_g_*as follows:

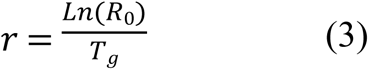

As *Ln (Ro)* remains approximately constant, it follows that a shorter generation time corresponds to a higher rate of increase. In the colder regions of Bolivia, *Tg* is nearly >2, while in the hotter areas, *Tg* is <0.5. With these contrasting values of 2 and 0.5, the rate of population growth in a hot region could be quadruple (*i.e.,* 2 / 0.5) that of a cooler region. Considering the representative localities, namely Camiri (hot Chaco region) and Cochabamba (cool Valleys region), where the mean number of generations per year stands at 1.73 and 0.86 respectively (as presented in Table 5), the generation time can be calculated as T*g* = 1 / 1.73 = 0.578 for Camiri and 1.163 for Cochabamba. Consequently, the rate of increase of *T. infestans* in Camiri is roughly *r* ≈ *Ln*(150) / 0.578 ≈ 8.67, which is twice that of Cochabamba (*r* ≈ 4.3).

Fig. 6 delineates the correlation between the extent of resistance observed across 21 localities and the number of generations per year. Notably, localities exhibiting elevated resistance levels (resistance ratio >10) are concurrently characterized by a heightened number of generations per year. Conversely, regions displaying lower resistance levels tend to manifest extended generation times, which aligns coherently with established theory [14].

**Fig. 6.**
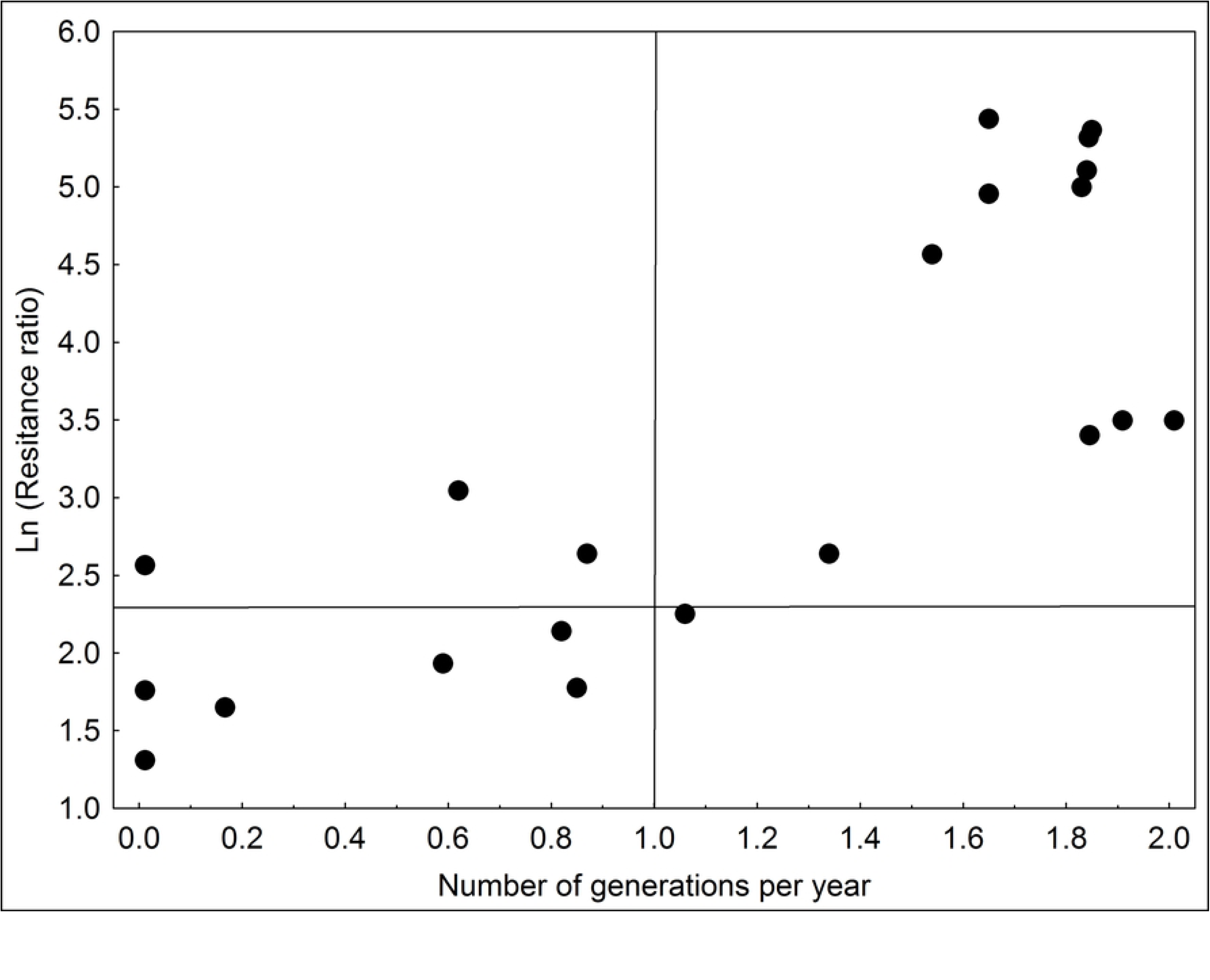
Relationship between the number of generations per year and deltamethrin resistance ratio (in Ln) for *T. infestans* strains collected in 21 localities of Bolivia. The horizontal line separates strains with low or no resistance (Resistance ratio *RR* <10, *i.e., Ln(RR*) <2.3) from strains with high levels of resistance in the upper part of the figure. The vertical line separates the univoltine (or less) *T. infestans* strains (number of generations per year ≤1) on the left side from almost bivoltine ones on the right side.

## Discussion

The study introduces a temperature-based developmental rate model for *T. infestans*, which holds universal applicability and can be readily employed in different contexts. The model unveiled that regions with higher temperatures experience an accelerated population growth. Significantly, these warmer regions with accelerated developmental rates also experience a higher number of generations per year. Additionally, there’s a discernible correlation between regions with elevated temperatures (thus fostering faster development and increased generational turnover) and higher levels of insecticide resistance.

The duration of the biological cycle of *T. infestans* has long captivated researchers’ attention. Initial laboratory investigations yielded cycle durations ranging from 220 to 240 days (temperatures unspecified) [16, 17], extending to 1.5 years [18]. Consequently, some authors inferred that under natural conditions, the cycle would endure for at least one year. Notably, an observation revealed a cycle duration range of approximately [84-134] days during winter and [84-102] days during summer, showcasing accelerated development within the 26-28°C range [19]. In Tucuman (Argentina), experiments conducted under insectary conditions mirroring the locale’s temperatures resulted in a complete cycle from egg to adults within three months. This led to the conclusion that *T. infestans* could potentially undergo two generations per year in this (hot) area, taking into account the deceleration of development in lower winter temperatures [20]. Further experiments conducted under controlled temperature settings reported cycle durations of 135 days at temperatures between 24-28°C [21], 134 days at 25°C, and 107 days at 33°C [22], as well as 123 days at 25°C and 152 days at 28°C [23], all aligning with the present findings. In a separate experiment, at a temperature of 26°C ±1°C, the mean duration from egg to adult was around 160 days [15], as compared to the approximately 140 days recorded in the current study at 25-27°C. In another study, the cycle duration at 37°C spanned [99-111] days [24], mirroring the present study’s results. Another study reported a cycle duration range of [69-109] days at 30°C and [132-327] days at 25°C [25]. Notably, while variations exist across different studies, these disparities likely stem from divergent environmental conditions, including insect crowding, feeding sources, feeding frequency, and other abiotic factors such as relative humidity. Nevertheless, overarching trends in the values remain comparable.

### Duration of incubation time

Our findings from controlled temperature chambers (Table 3) align harmoniously with prior research. Other investigations affirm that the average incubation period for the egg-stage was roughly 20 days at 26°C [15], 27.4 days at 25°C, and 14.4 days at 30°C [25], as well as 20-24 days at 25°C and 11-13 days at 33°C [22], with another study recording 18.9 ± 1.6 days at 25-27°C [19]. Earlier studies, albeit not specifying experimental temperatures, cited an average within the range of 20-25 days [16], while in Rio de Janeiro (Brazil) an experiment indicated 20.2 ± 2.4 days [19]. Longer incubation periods, extending to 46 days, were also documented [22], and in Cochabamba, Bolivia, where ambient temperatures tend to be lower, the estimated incubation time was 35 days [26].

### Duration of nymphal stages

Once again, our findings (Table 3) resoundingly align with previous experimental results. In parallel investigations, the total duration of the entire nymphal stage was assessed at 141 days at 26°C [15] or 102.4 ±10.9 days at the same temperature [19]. In contrast, our study indicates that the complete nymphal stage duration (from N1 to N5) spans approximately 100-120 days, contingent on these same temperature conditions.

As delineated in Table 3, the mean duration of each stage (N1 to N4) at various temperatures (excluding 20°C) exhibits only marginal variability, amounting to a global mean duration of approximately 17 days. The global median duration equals 17 days, featuring an interquartile range of approximately [15 - 19] days. Notably, the N5 stage presents a mean duration nearly twice as long (global mean ≈ 37 days), mirroring patterns observed in other research studies [15]. The protracted duration of the N5 stage could potentially signify a preliminary indication of diapause [27], offering a potential adaptive strategy in response to environmental fluctuations [28].

### The optimal temperature

The Briere-1 model’s computation of the optimal temperature at 33.5°C for the entire cycle is in congruence with the suggested optimal rearing temperature of 30°C for the majority of Triatominae species. Across various development stages, the optimal temperature values remained consistently close, falling within the range of [33.1 – 34.5°C] (Table 4).

### The development threshold temperature

The natural development of *T. infestans* exhibits a stronger dependence on the minimal temperature rather than the average temperature [29, 30], thereby highlighting the significant role of the development threshold temperature. Our current findings established this threshold at 15.7°C, encompassing a range of [14.2 – 17.7°C] across various development stages (Table 4). This is in harmony with the 16°C threshold reported in a previous study [29], further supported by the observed absence of molting at 16°C in our study. While *T. infestans* can endure lower temperatures, their feeding activity halts at 10°C [27]. The temperature range conducive to normal *T. infestans* development has been estimated to be above 20°C. This species has been characterized as temperate, displaying adaptability to regions characterized by wide temperature fluctuations. This adaptability is mirrored by the thermal breadths calculated by the Briere-1 model in our study.

### The upper lethal temperature

An upper limit of 37°C has been proposed for the rearing of *T. infestans* [24]. This aligns with our experimental findings at 37°C, which unveiled a noticeable decline in the development rate of *T. infestans*. This decline implies a close approximation to the critical (lethal) temperature threshold, where survival becomes compromised. In fact, the Briere-1 model’s estimation positioned the upper lethal temperature at 39.3°C, encompassing a range of [38.9 – 40.1°C] across various development stages (Table 4).

### The number of generations per year

Across previous studies, the span from one egg-laying to another has been estimated at 143 days [19] and 163 days at 26°C [15], equating to approximately 2.55 and 2.24 generations per year, respectively. These outcomes harmonize with the computed number of generations per year for the four representative localities as detailed in our present study (Table 5). Notably, a robust linear relationship exists between the number of generations per year within these localities and the mean annual temperature (the equation is *Number of generations = −1.8131 + 0.1562 * Temperature, R^2^* = 0.99). Additionally, the Rabinovich [15] data align precisely with the extension of the regression line (Fig. 7).

**Fig. 7.**
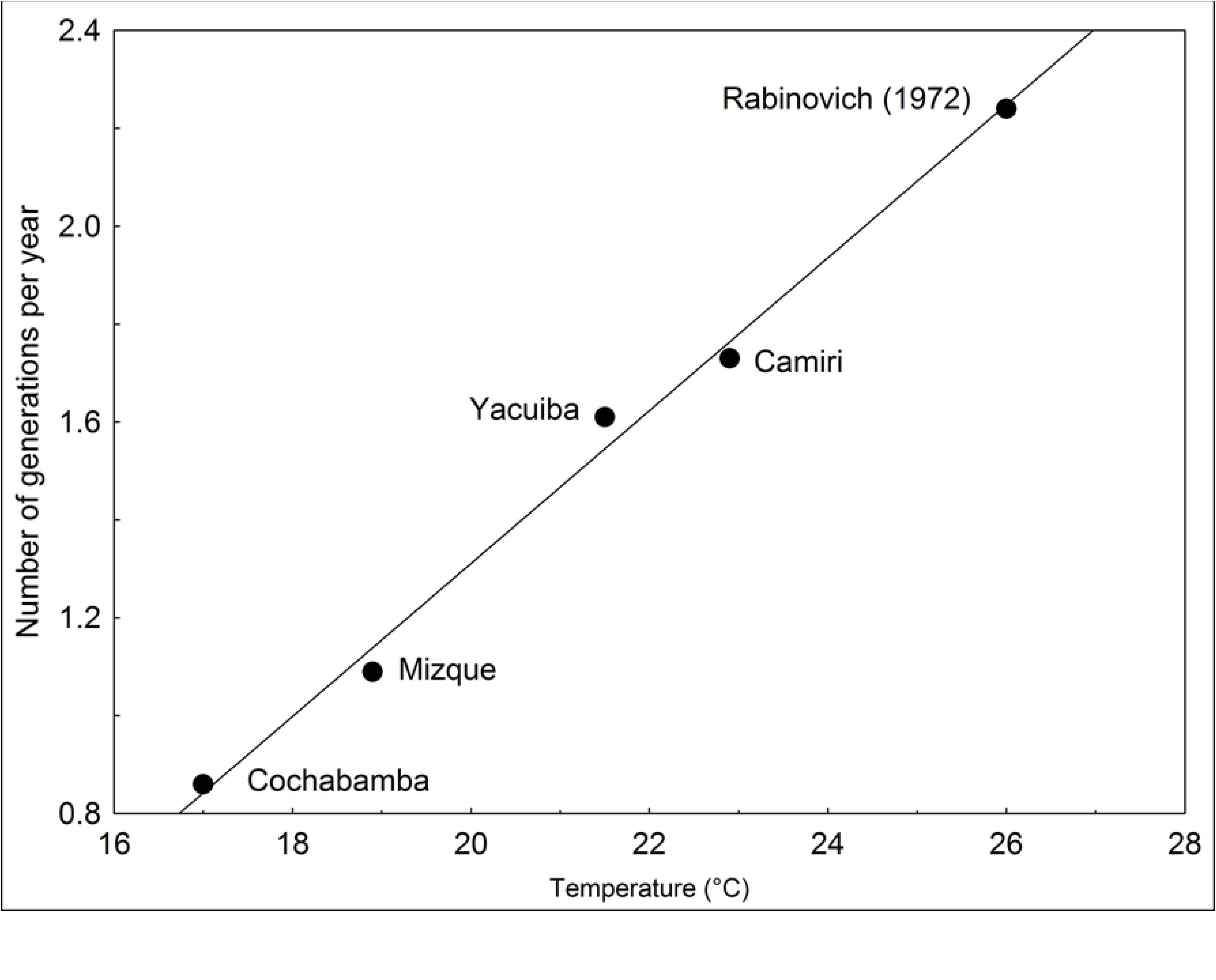
Relationship between the mean annual temperature (°C) of the representative localities and the mean number of generations per year. The Rabinovich (1972) data is projected as supplementary data of the regression line.

Within the geographic range of *T. infestans*, the number of generations per year, and consequently the rate of population expansion, can differ by a factor of up to two, and potentially even four or more, between the cooler Andean Valleys and the hotter low-lying regions. As a result, the population dynamics of this vector unfold more swiftly in such warmer areas, leading to a hastened emergence of resistance to insecticides. The emphasis on generation time as a factor influencing the efficacy of chemical control efforts partially elucidates the scenario wherein Bolivia managed to attain vector transmission-free certification in 23 out of 154 endemic municipalities located in the cold areas of the La Paz and Potosi Department in 2010 and 2012. In contrast, in the hotter regions, control efforts have proved and continue to pose significant challenges.

### Other factors

In the scope of our current study, the experimental conditions within the climatic chambers were optimal, ensuring triatomines had sufficient food availability (once a week) and stable relative humidity. Consequently, the phenological model delineates optimal development, where variability is likely attenuated compared to natural conditions, predominantly arising from temperature fluctuations. It is essential to note that our model’s results are rooted in mean daily temperatures, differing from the dynamic daily oscillation between a minimum and maximum temperature. Nonetheless, our findings uphold alignment with other studies conducted in natural settings. Thus, while the present phenological model remains simplistic, it appears adequate for (1) accurately depicting *T. infestans’* development in response to temperature and (2) facilitating comparisons between different regions and localities. The overarching insights from the model, particularly regarding distinctions in regional “generation times influenced by temperature” and the “emergence of insecticide resistance linked to the number of generations per year”, exhibit consistency and coherence.

## Author contributions

**Conceptualization:** Frédéric Lardeux

**Data curation:** Frédéric Lardeux

**Formal Analysis:** Frédéric Lardeux

**Funding acquisition:** Frédéric Lardeux

**Investigation:** Stéphanie Depickère, Frédéric Lardeux

**Methodology**: Frédéric Lardeux

**Resources**: Stéphanie Depickère, Rosenka Tejerina

**Validation**: Stéphanie Depickère, Rosenka Tejerina

**Project administration:** Stéphanie Depickère

**Writing – original draft**: Frédéric Lardeux, Stéphanie Depickère

**Writing – review & editing:** Frédéric Lardeux, Rosenka Tejerina

## Acknowledgements

To Hubert Mazurek (IRD, UMR LPED) for his advices in GIS manipulation. To Alberto Llanos and Guillermo Aliaga for the help in *T. infestans* colony maintenance. To the Abdus Salam International Centre for Theoretical Physics of Trieste, Italy (ICTP) for supporting S. Depickère.

## Supporting Information Captions

The supporting information files are deposited in the Harward Dataverse (https://dataverse.harvard.edu/) with dataset DOI: 10.7910/DVN/Y0KT1L and will stay unpublished until manuscript revision and acceptation.

**S1 Table. The 7988 individual development rates**. Measures of duration of each stage, from egg to N1, N1 to N2, N2 to N3, N3 to N4, and N4 to N5, when insects are exposed to six different constant temperatures.

**S2 Table. Simulation of the development cycle of *T. infestans***. Simulations of the whole development cycle of *T. infestans* with the @RISK software, at different constant temperatures.

**S3 Table. Mean daily temperatures from 2012 to 2017**. Mean daily temperatures in the four representatives cities selected in the present study.

## References

1. Lardeux F, Depickère S, Aliaga C, Chavez T, Zambrana L. Experimental control of *Triatoma infestans* in poor rural villages of Bolivia through community participation. Trans R Soc Trop Med Hyg. 2015;109(2):150–8. doi: 10.1093/trstmh/tru205. PubMed PMID: WOS:000350102900010.

2. Hopkins T, Goncalves R, Mamani J, Courtenay O, Bern C. Chagas disease in the Bolivian Chaco: Persistent transmission indicated by childhood seroscreening study. Int J Infect Dis. 2019;86:175–7. doi: 10.1016/j.ijid.2019.07.020. PubMed PMID: 31357060.

3. Lardeux F, Depickère S, Duchon S, Chavez T. Insecticide resistance of Triatoma infestans (Hemiptera, Reduviidae) vector of Chagas disease in Bolivia. Trop Med Int Health. 2010;15(9):1037–48. doi: DOI 10.1111/j.1365-3156.2010.02573.x. PubMed PMID: WOS:000280662700008.

4. Depickère S, Buitrago R, Siñani E, Baune M, Monje M, López R, et al. Susceptibility and resistance to deltamethrin of wild and domestic populations of *Triatoma infestans* (Reduviidae: Triatominae) in Bolivia: new discoveries. Mem Inst Oswaldo Cruz. 2012;107(8):1042–7. PubMed PMID: WOS:000328105100013.

5. Gürtler RE, Canale DM, Spillmann C, Stariolo R, Salomon OD, Blanco S, et al. Effectiveness of residual spraying of peridomestic ecotopes with deltamethrin and permethrin on *Triatoma infestans* in rural western Argentina: a district-wide randomized trial. Bull WHO. 2004;82(3):196–205. PubMed PMID: 15112008.

6. Gorla DE, Ortiz RV, Catala SS. Control of rural house infestation by *Triatoma infestans* in the Bolivian Chaco using a microencapsulated insecticide formulation. Parasites & Vectors. 2015;8. doi: 10.1186/s13071-015-0762-0. PubMed PMID: WOS:000353946400001.

7. Chuine I, Regniere J. Process-based models of phenology for plants and animals. Annu Rev Ecol, Evol Syst. 2017;48:159–82. doi: 10.1146/annurev-ecolsys-110316-022706. PubMed PMID: WOS:000415250000008.

8. Picollo MI, Wood E, Zerba E, Licastro SA, Ruveda MA. Metodos de laboratorio para medir la actividad de insecticidas en *Triatoma infestans*. Acta Bioq Clín Latinoam. 1976;10:67–71.

9. Lardeux FJ, Tejerina RH, Quispe V, Chávez TK. A physiological time analysis of the duration of the gonotrophic cycle of *Anopheles pseudopunctipennis* and its implications for malaria transmission in Bolivia. Malaria Journal. 2008;7. doi: 10.1186/1475-2875-7-141. PubMed PMID: WOS:000258435900001.

10. Mirhosseini MA, Fathipour Y, Reddy GVP. Arthropod development’s response to temperature: a review and new software for modeling. Ann Entomol Soc Am. 2017;110(6):507–20. doi: 10.1093/aesa/sax071. PubMed PMID: WOS:000414410300001.

11. Brière JF, Pracros P, Le Roux AY, Pierre JS. A novel rate model of temperature-dependant development for arthropods. Environ Entomol. 1999;28:22–9.

12. Rebaudo F, Struelens Q, Dangles O. Modelling temperature-dependent development rate and phenology in arthropods: The DEVRATE package for R. Methods Ecol Evol. 2018;9(4):1144–50. doi: 10.1111/2041-210x.12935. PubMed PMID: WOS:000429421800031.

13. Rojas M, Avalos M, Rocha V, Gorla D. Distribución biogeográfica de los triatominos en Bolivia. Discriminación de la distribución de las especies en relación a variables ambientales. In: Rojas M, editor. Triatominos de Bolivia y la enfermedad de Chagas. La Paz, Bolivia: Ministerio de Salud y Deportes; 2007. p. 69–137.

14. Anderson RM, May RM. Infectious diseases of humans: dynamics and control. Oxford and New York: Oxford University Press; 1991. 123 p.

15. Rabinovich JE. Vital statistics of Triatominae (Hemiptera: Reduviidae) under laboratory conditions. I. Triatoma infestans Klug. J Med Entomol. 1972;9(4):351–70.

16. Neiva A. Informaçoes sobre biologia da Vinchuca, *Triatoma infestans* Klug. Mem Inst Oswaldo Cruz. 1913;5:24–31.

17. Abalos JW, Wygodzinsky PW. Las Triatominae Argentinas (Reduvidae, Hemiptera). Publ Inst Med Reg 1951;601(2):1–179.

18. Maggio C, Rosenbusch F. Studien über die Chagas Krankheit in Argentinien und die Trypanosomen der Vinchucas. Centralbl f Bakt Orig. 1915;77:40.

19. Perlowagora-Szumlewicz A. Estudos sôbre la biologia do Triatoma infestans, o principal vetor da doença de Chagas no Brasil. (Importância de algumas de suas caracteristicas biologicas no plenejamento de esquemas de combate a esse vetor). Rev Bras Malariol D Trop. 1969;21(1):117–59.

20. Romaña C. La enfermedad de Chagas y otras tripanosomiasis humanas en Americas. 5^to^ Cong Int Med Trop Malaria. 1953;1:187–201.

21. Perlowagora-Szumlewicz A. Laboratories colonies of Triatominae, biology and population dynamics. PAHO Sci. 1975;318:63–82.

22. Hack WH. Estudios sobre biologia del *Triatoma infestans* (Klug, 1834) (Hemiptera, Reduviidae). An Inst Med Reg. 1955 4:125–47.

23. Correa FMA. Estudo comparativo do ciclo evolutivo do Triatoma infestans alimentado em diferentes animáis — (Hemiptera, Reduviidae). Pap Avuls Zool. 1962;15:177–200.

24. Pessoa SB, Barros NV. Criaçao do *Triatoma infestans* na temperatura de estufa. Fôlha Med. 1939;20:285–7.

25. Juárez M. Comportamento do *Triatoma infestans* sob várias condições de laboratório Rev Saúde Públ. 1970;4:147–66.

26. Borda PMR. Algunos nuevos aspectos sobre biología y ecología de Triatoma infestans Klug, 1834, y su enemigo natural Telenomus fariai Lima, 1927. Breves notas referentes a Trypanosoma cruzi Chagas, 1909. Rev Per Entomol. 1971;14(2):379–85. PubMed PMID: 19750521935.

27. Joërg ME. Influencia de temperaturas fijas en periodos anuales sobre metamorfosis y fertilidad de *Triatoma infestans*. Bol Chil Parasitol. 1962;17:17–9.

28. Menu F, Ginoux M, Rajon E, Lazzari CR, Rabinovich JE. Adaptive developmental delay in Chagas disease vectors: an evolutionary ecology approach. PLoS Neglected Tropical Diseases. 2010;4(5). doi: 10.1371/journal.pntd.0000691. PubMed PMID: ISI:000278601000021.

29. Gorla DE, Schofield CJ. Population dynamics of *Triatoma infestans* under natural climatic conditions in the Argentine Chaco. Med Vet Entomol. 1989;3(2):179–94. PubMed PMID: 2519662.

30. Medone P, Ceccarelli S, Parham PE, Figuera A, Rabinovich JE. The impact of climate change on the geographical distribution of two vectors of Chagas disease: implications for the force of infection. Philos Trans R Soc B-Biol Sci. 2015;370(1665). doi: 10.1098/rstb.2013.0560. PubMed PMID: WOS:000350829800008.

